# Eye Movements in Silent Visual Speech track Unheard Acoustic Signals and Relate to Hearing Experience

**DOI:** 10.1101/2024.09.10.610824

**Authors:** Kaja Rosa Benz, Anne Hauswald, Nina Suess, Quirin Gehmacher, Gianpaolo Demarchi, Fabian Schmidt, Gudrun Herzog, Sebastian Rösch, Nathan Weisz

**Affiliations:** Centre for Cognitive Neuroscience, Department of Psychology, Paris-Lodron-University of Salzburg, Salzburg, Austria; Neuroscience Institute, Christian Doppler University Hospital, Paracelsus Medical University Salzburg, Salzburg, Austria; Department of Experimental Psychology, University College London, United Kingdom; Wellcome Centre for Human Neuroimaging, University College London, United Kingdom; Deaf Outpatient Clinic, University Hospital Salzburg (SALK), Salzburg, Austria; Clinic and Polyclinic for Otorhinolaryngology, University Hospital Regensburg, Regensburg, Germany

## Abstract

Behavioral and neuroscientific studies have shown that watching a speaker’s lip movements aids speech comprehension. Intriguingly, even when videos of speakers are presented silently, various cortical regions track auditory features, such as the envelope. Recently, we demonstrated that eye movements track low-level acoustic information when attentively listening to speech. In this study we investigated whether ocular speech tracking occurs during visual speech and how it influences cortical silent speech tracking. Furthermore, we compared the data of hearing individuals with congenitally deaf individuals, and those with acquired deafness or hearing loss (DHH; Deaf or hard of hearing) to assess how auditory deprivation (early vs. late onset) affects neural and ocular speech tracking during silent lip-reading. Using magnetoencephalography (MEG), we examined ocular and neural speech tracking of 75 participants observing silent videos of a speaker played forward and backward. Our main finding is a clear ocular unheard speech tracking effect with a dominance <1 Hz, which was not present for the lip movements. Similarly, we observed a <=1.3 Hz effect of neural unheard speech tracking in temporal regions for hearing participants. Importantly, neural tracking was not directly linked to ocular tracking in this study. Strikingly, across different listening groups, deaf participants with auditory listening experience showed higher ocular speech tracking than hearing participants, while no ocular speech tracking effect was revealed for congenitally deaf participants in a very small sample. This study extends our previous work by demonstrating the involvement of eye movements in speech processing, even in the absence of acoustic input.

## 1. Introduction

Successful speech perception in naturalistic settings normally requires the joint activity of multiple sensory modalities, integrated into a coherent experience. In hearing individuals, auditory speech is integrated with the visually observed movements of the speaker. Meanwhile, deaf individuals without cochlear implants, who process verbally communicated speech, can only exploit unimodal visual information, primarily encoded in the mouth movements of the observed speaker.

To successfully process auditory speech, neural activity in relevant processing regions needs to temporally align its excitability phases with speech features (e.g., syllables;(Doelling et al., 2014)). This concept, known as neural speech tracking (Obleser & Kayser, 2019), is most commonly operationalized by quantifying the relationship between the time series of neural activity and the envelope of the speech signal. Neural speech tracking occurs passively and is further enhanced when attention is directed to the speech (Vanthornhout et al., 2019).

When congruent lip movements are perceived in addition to auditory speech, the lip movements support understanding of degraded (e.g., noisy, vocoded) speech in hearing individuals (Haider et al., 2022, 2024; Macleod & Summerfield, 1987; O’Sullivan et al., 2020; Ross et al., 2006). A possible explanation is that processing a speaker’s lip movements enhances speech tracking (Crosse et al., 2015, 2016). Evidence for this has been found and shown to be especially pronounced in challenging listening situations (Haider et al., 2022). When predicting EEG signals at posterior electrodes of subjects who watched videos of silent speech, adding the unheard speech envelope significantly enhanced accuracies (O’Sullivan et al., 2017). Going beyond this finding, we could show that cortical tracking of the unheard speech envelope during silent videos was linked to the intelligibility of auditory speech: this effect was particularly pronounced in occipital regions, with higher speech-brain coherence for forward versus reversed videos (Hauswald et al., 2018; Suess et al., 2022). Importantly, when using visual speech information, no difference is observed between forward versus backward presented videos, underlining that this effect is specific to the corresponding but unheard auditory information. As this phenomenon is found in passive settings (Hauswald et al., 2018; Suess et al., 2022),it suggests that speech features are more or less automatically activated from purely visual input. The processes contributing to this phenomenon have, however, remained elusive so far.

A possible contributor could be ocular speech tracking, the phenomenon where eye movements track auditory speech (Gehmacher et al., 2024). This finding extends Jin et al., (2018), who observed that ocular muscle activity is synchronized to artificially rhythmic continuous speech. Importantly, Gehmacher et al., (2024) showed that ocular speech tracking was modulated by attention not only across sensory modalities but also within the auditory modality in a multispeaker situation (for an extended replication, see Schubert et al., 2023). Furthermore, neural speech tracking significantly decreased when eye movements were taken into account (Gehmacher et al., 2024). These findings suggest that ocular speech tracking might help the listener track relevant acoustic input, especially when understanding becomes more challenging. We hypothesize that if eye movements are generally involved in the processing of speech, then ocular speech tracking should also, in line with neural speech tracking, be observed during silent speech.

Furthermore, this process should critically depend on the learned association between processing visual cues of a speaker and the uttered auditory speech. Therefore, in the present study we also investigate how the visuophonological transformation (ocular and neural tracking of unheard speech) differs depending on audiovisual listening experience. Congenitally deaf individuals are born without hearing, whereas the deaf participants with listening experience lost their hearing after being exposed to audio-visual speech. Including congenitally deaf, late DHH (Deaf or hard of hearing), and hearing individuals, our sample includes three groups with different experiences with spoken language. Auditory sensory deprivation combined with the necessity to rely on visual signals can give us important insights into multisensory speech processing. In the deaf community, ears aren’t pivotal for speech perception, whereas the eyes may play an even more significant role. If ocular and neural speech tracking during silent lip reading depends on the learned audiovisual experience concerning critical periods of spoken language development, we predict that resulting patterns should be more similar between a hearing and a deaf sample with audiovisual listening experience, while the congenitally deaf group should show deviating patterns.

Using estimations of eye movements from magnetoencephalographic data, and coherence as a measure of speech tracking, we show that in hearing participants, eye movements track silent speech in a low (∼1 Hz) frequency range, while we do not find this effect for the visual information (lip movements). In the same frequency range, we also observe a cortical network of auditory and motor regions with analogous neural tracking effects. Interestingly, controlling for ocular speech tracking does not reduce neural speech tracking effects in this frequency range. Considering listening experience, we found enhanced ocular unheard speech tracking for the acquired DHH group compared to the hearing group. Crucially, we found no ocular unheard speech tracking in the congenitally deaf group, underlining that this effect is experience dependent.

## 2. Results

We investigated whether ocular tracking of unheard speech exists in hearing subjects using coherence. Therefore, we compared speech tracking in a forward vs. backward condition (Fig. 1A). Subsequently, we examined the relationship of ocular speech tracking to neural speech tracking using partial coherence. In a second step, we compared ocular and neural speech tracking during silent lip-reading across three distinct groups: a hearing group, a group of deaf individuals without prior exposure to auditory language, and a group of deaf individuals with prior experience in auditory language. As a measure for eye movements, we used the strongest and most reliable ocular ICA component, mostly capturing vertical eye movements (including the blinks) as a substitute for EOG/Eye-tracking allowing consistently good data quality over all subjects and the investigation of the full sample.

**Fig. 1:**
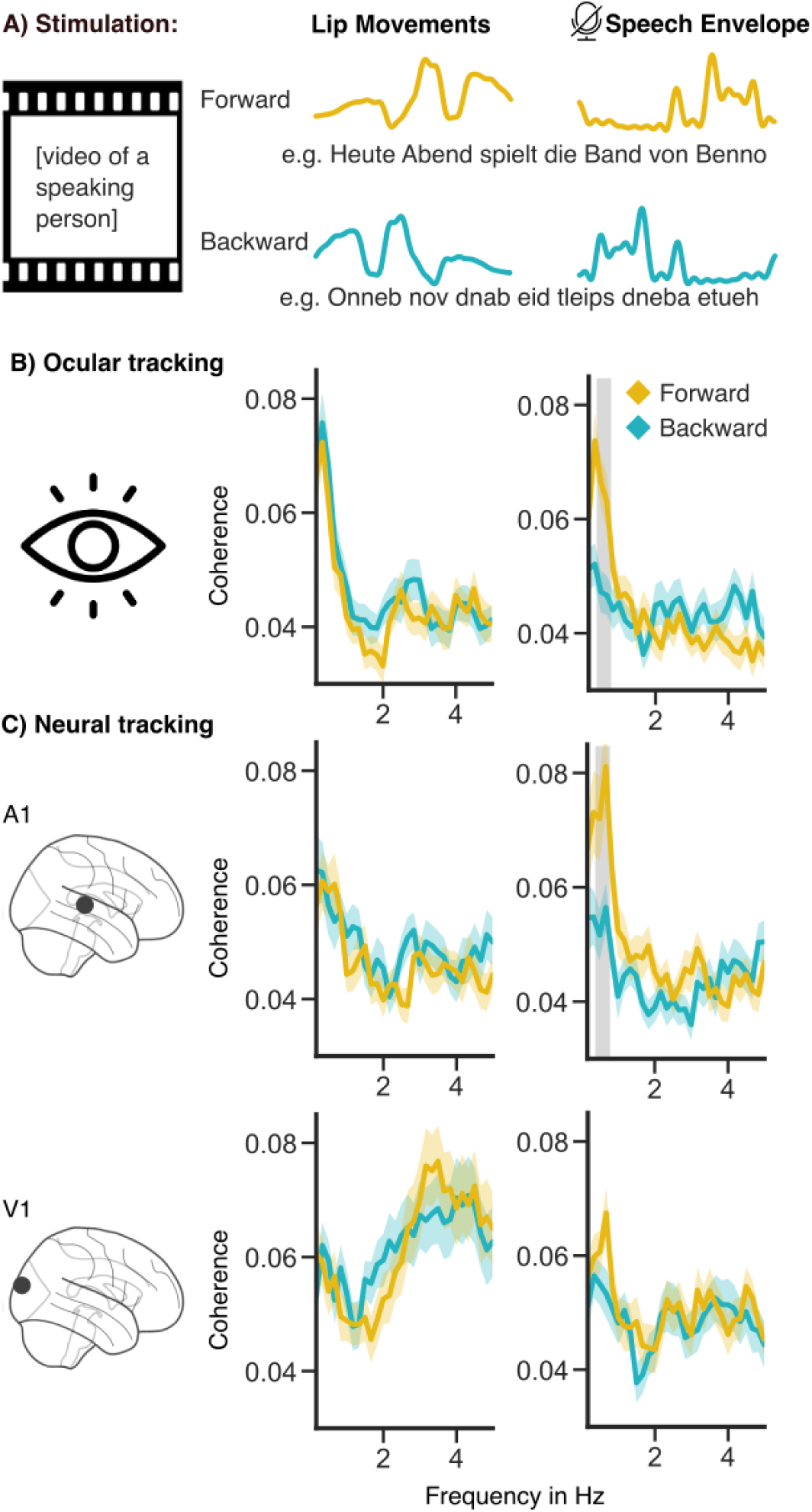
A) Stimulus Material: Participants observed videos of lip movements played either forward or backward. Lip movements (opening) and the corresponding unheard speech envelope were extracted as a continuous signal; B) Analysis of the hearing group: Results of coherence calculation between the speech envelope and lip movements using the selected ICA eye component. C) Coherence calculation at the strongest voxel in the primary auditory and visual cortex. Significant frequency clusters are marked in gray (N=49; p < 0.01). The face of the speaking person is included in accordance with bioRxiv guidelines. See supplements for the same figure for the deaf groups.

### 2.1. Ocular tracking of silent speech in the hearing group

We first addressed the question of whether speech tracking is present in the ocular data. Participants watched silent videos of speakers in both forward and reverse directions. Coherence between eye movements and the lip movements or the unheard speech envelope of the corresponding video was calculated in the frequency range between 0.16 and 5 Hz. This range was chosen because speech tracking (Chalas et al., 2023; Schmidt et al., 2023) and silent speech tracking (Bourguignon et al., 2020; Suess et al., 2022) have been recently found to be most pronounced in the low-frequency (delta) range.

Comparing ocular speech tracking in the forward versus backward conditions, we found a cluster in the frequency range from 0.33 to 0.83 Hz (*p < 0.01*, Fig. 1B) with increased tracking in the forward condition. Since acoustic and visual information are highly correlated, it is necessary to test whether tracking of visual information is enhanced in the same frequency range. For lip movements, no cluster was revealed, emphasizing that ocular speech-tracking effects are specific to unheard auditory features. Overall, this analysis illustrates that the effects of transforming speech from a purely visual to an auditory format can also be captured via eye movements.

### 2.2. Cortical tracking of silent speech in the hearing group

As we wanted to investigate the relationship between the previously established neural effects (Aller et al., 2022; Bourguignon et al., 2020; Bröhl et al., 2022; Hauswald et al., 2018; Suess et al., 2022) and the ocular effects presented here, it was crucial to first replicate the neural effects for relevant ROIs and identify further cortical regions at a whole-brain level. For this purpose, we conducted a source analysis of the MEG data and calculated coherence (range: 0.16 to 5 Hz) between each voxel and the lip movements, as well as the unheard speech envelope. To compare the ocular coherence spectra with the neural frequency spectra of ROIs, we extracted the source with the strongest effect within the bilateral primary visual and primary auditory cortex (Fig. 1C). This showed strongly overlapping patterns with the ocular speech tracking effect (Fig. 1B), and in the primary auditory cortex, we also found an effect of enhanced speech tracking from 0.33 to 0.83 Hz (p < 0.01), while in the visual cortex no cluster appeared Examining this phenomenon on a whole-brain level, we observed that the strongest tracking of lip movements occurred at 4 Hz in occipital regions (see Suppl. Mat.). Since the speech envelope correlates with lip movements, we initially examined whether there were any distinctions in lip tracking between the forward and backward conditions. In the cluster permutation t-test over the whole brain, no clusters were revealed. Subsequently, we tested for differences in speech tracking in the forward vs. backward condition for the whole brain. The data-driven cluster included the primary auditory cortex, somatosensory and – motor areas, the inferior frontal gyrus and the temporoparietal junction (0.16–0.83 Hz, p < 0.01; Fig. 2A). Overall, these results illustrate a network of especially temporal and motor brain regions for tracking the speech envelope during silent videos in a frequency range very similar to the ocular speech tracking effect.

**Fig. 2:**
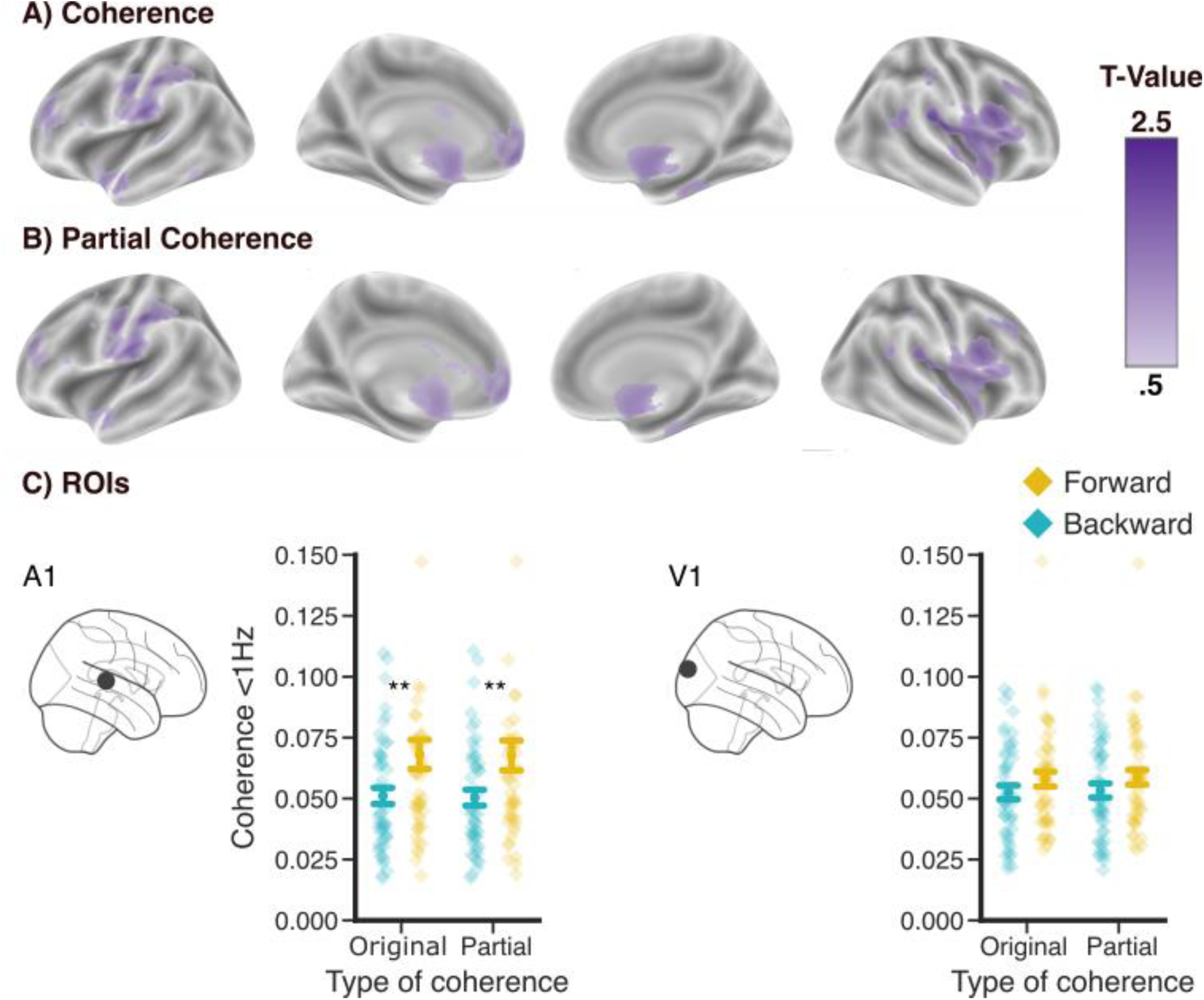
Role of eye movements in the cortical effects in the hearing group. A) Significant clusters of the forward vs. backward comparison of the speech-brain coherence. B) Significant clusters of the forward vs. backward comparison of the speech-brain coherence remain the same after controlling for eye movements. C) Coherence <1 Hz in the A1 and the V1 for coherence and partial coherence. Cluster permutation threshold was a p-value of 0.01. Error bands reflect the standard error (N=49). Asterisks indicate levels of significance: ** = p < 0.01.

### 2.3. Cortical speech tracking controlled for ocular speech tracking in the hearing group

It has been shown that ocular and neural activity share contributions to speech processing (Gehmacher et al., 2024). To quantitatively test this contribution to the tracking of unheard speech, we used partial coherence to control for the influence of ocular activity on speech-brain coherence. This was followed by the same analysis as described above, comparing the forward vs. backward condition using a whole-brain cluster-based permutation test. Even after applying partial coherence, a significant neural speech tracking effect was observed from 0.16 to 1.33 Hz (p < 0.01), with a highly overlapping spatial distribution (Fig. 2B). Comparing the coherence results, including the eye movements, with the partial-coherence results controlled for the eye movements, a cluster-based permutation t-test over the whole brain did not reveal any significant differences.

For illustration purposes, the voxel revealing the strongest effects in the primary auditory and primary visual cortex (same voxel as in Fig. 1C) was selected and averaged over frequencies below 1 Hz (Fig. 2C). In temporal regions, we found an effect of condition (F(48,1) = 11.58, p = 0.001), which was not observed occipitally (F(48,1) = 2.43, p = 0.12). However, we did not observe a significant difference between coherence and partial coherence. Based on these results, ocular activity—at least when captured using the ICA-eye component - does not appear to drive the speech-brain coherence effects.

### 2.4. Role of hearing experience in neural and ocular silent speech tracking

After establishing that hearing individuals track unheard speech at slow frequencies while watching silent videos of speakers, we explored how this pattern varies with individuals’ audiovisual spoken language experience. Specifically, we were interested in whether deaf individuals, who were born deaf or had some experience with spoken language, exhibit different patterns. We conducted the same analysis for the acquired DHH group (N = 18) as we did for the hearing group and found that the acquired DHH group also exhibited the ocular speech tracking effect (0.16-0.66 Hz, p < 0.01). Additionally, when focusing on the temporal regions of interest, we found a neural effect of speech tracking (0.16-0.83 Hz, p < 0.01). Importantly, we found that this neural effect could not be explained by ocular tracking. However, on a whole-brain level, these effects were not as pronounced as in the hearing group, which may be attributed to the smaller sample size in the acquired DHH group (see supplementals). For the congenitally deaf group (N = 7), we did not find any significant effects for either ocular or neural speech tracking.

To test for differences in ocular speech tracking between the three groups of hearing experience, we first calculated the difference between the forward and backward condition and used it as the dependent variable in a 2×3 ANOVA (modality: mouth area, speech envelope * group; Fig. 3A). The two-way ANOVA revealed a significant effect of group (F(72,2) = 4.97, p < 0.01), modality (F(72,1) = 7.73, p < 0.01), and also an interaction (F(72,2) = 3.1, p = 0.05). Post hoc tests revealed that the forward vs. backward difference for the speech envelope is higher than for the lip movements across all groups (T(74) = -2.7, p < 0.01). Moreover, the acquired DHH group shows significantly higher ocular tracking than the hearing group (T(21.98) = 2.05, p = 0.05) and the congenitally deaf group (T(22.9) = 2.74, p = 0.01). The interaction effects show that the acquired DHH group exhibits significantly higher ocular lip tracking in the forward compared to the backward condition than the hearing group (T(21.51) = 2.59, p < 0.05) and the congenitally deaf group (T(23.99) = 2.86, p < 0.01).

**Fig. 3:**
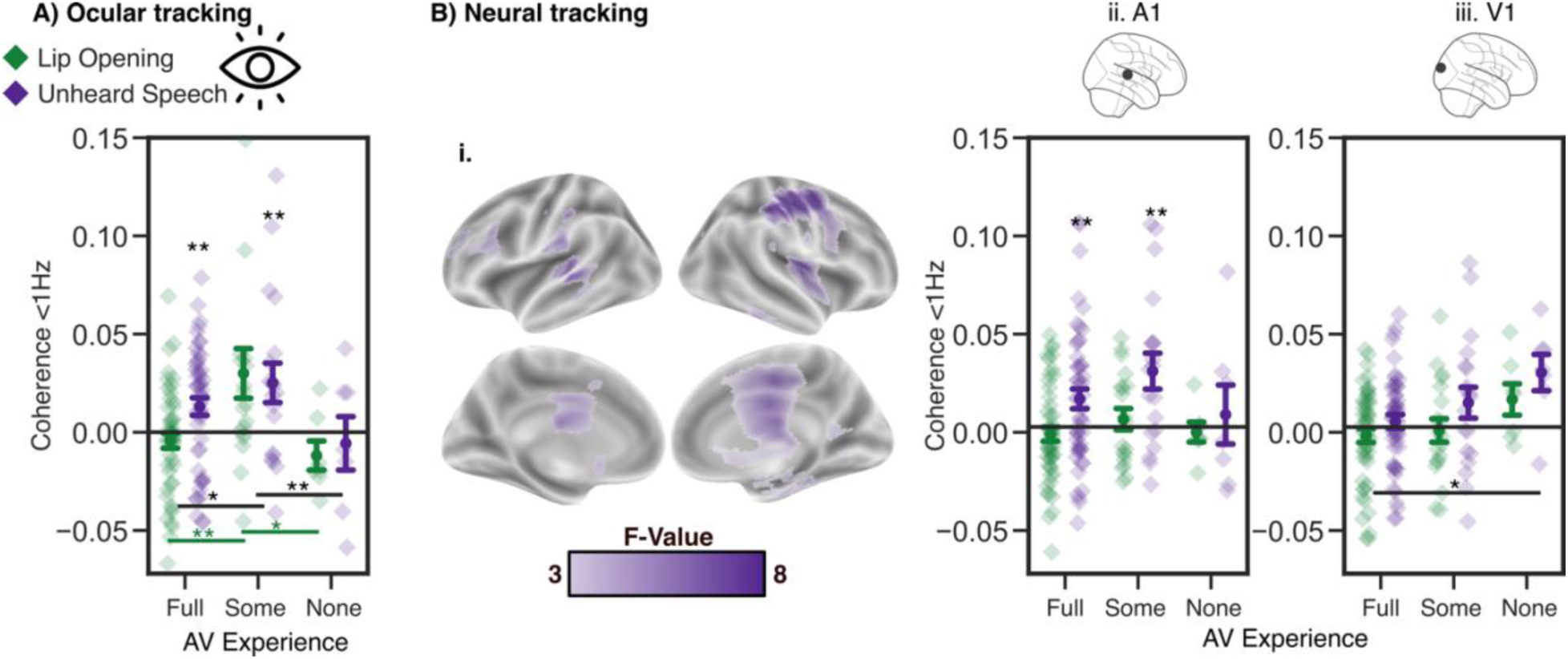
A) Ocular tracking: <1 Hz Forward - Backward tracking of the unheard speech envelope for the three hearing groups. The hearing and the DHH group (with audiovisual listening experience) show the ocular unheard speech tracking effect, which is not the case for the congenitally deaf group (see stars above the jitter plot). Furthermore, the DHH group shows increased tracking in the forward compared to the backward condition in general (see lines in black), and increased lip tracking in the forward condition (see lines in green) compared to the other groups. B) i. The main effect across all subjects showed higher forward vs. backward differences in speech than in lip tracking. ii. A1: <1 Hz tracking in the auditory cortex: The hearing and the DHH groups show the unheard speech tracking effect. iii. V1: <1 Hz tracking in the visual cortex: The congenitally deaf group has higher forward vs. backward tracking than the hearing group. The same voxels were selected as in *Figs. 1 & 2*. For the whole-brain ANOVA, the cluster permutation threshold was a p-value of 0.01. Error bands reflect the standard error. Asterisks indicate levels of significance: * = p < 0.05, ** = p < 0.01.

Hence, across all groups, the speech envelope is tracked more strongly in the forward condition, which is not the case for the lip movements. Furthermore, the acquired DHH group showed the highest ocular tracking and increased tracking of the lip movements compared to the other groups. The differences in ocular unheard speech tracking between the three groups with different audiovisual language experiences imply that ocular unheard speech tracking is relevant in audiovisual speech processing.

To test for differences on a neural level between the three groups of hearing experience, we conducted the same 2×3 ANOVA as for the ocular speech tracking, with cluster permutation over the whole brain. This resulted in a main effect showing a higher speech-tracking difference in the forward vs. backward condition than a lip-tracking difference (0.16-1.66 Hz, p < 0.001; Fig. 2B.i). However, no main effect of group nor an interaction was found. Focusing on the regions of interest, the same main effect of modality was reflected in A1 (F(72,1) = 19.55, p < 0.001; Fig. 2B.ii) and V1 (F(72,1) = 7.09, p < 0.01; Fig. 2B.iii). Furthermore, in V1, there was also a group effect present (F(72,2) = 3.5, p < 0.05), showing increased forward tracking for the congenitally deaf group compared to the hearing group (post hoc: T(7.68) = 2.6, p < 0.05).

Taken together, on a cortical level, the hearing and acquired DHH groups reveal similar patterns; both track the unheard speech envelope more strongly in the forward condition with the eyes and the A1. This is not the case for the congenitally deaf group, which does not show those patterns but shows increased occipital tracking compared to the hearing group.

## 3. Discussion

In the present study, we compared how eye movements track unheard speech between groups with different levels of audiovisual listening experience. While previous studies have explored ocular speech tracking (Gehmacher et al., 2024; Schubert et al., 2023), we aimed to establish a direct link between ocular unheard speech tracking, neural unheard speech tracking, and audiovisual listening experience. Overall, the results show stronger ocular unheard speech tracking for forward-played videos compared to reversed ones, indicating that the eyes track meaningful speech from silent lip movements. On a neural level, this effect was primarily observed in temporal regions and persisted even after controlling for eye movements. Among the deaf groups, congenitally deaf participants did not show any significant tracking effects, whereas the acquired DHH group exhibited stronger ocular speech tracking than the other groups. Neurally, while the hearing and DHH groups mainly tracked speech temporally, tracking of both lip opening and unheard speech tracking in the congenitally deaf group was enhanced in occipital regions.

### Eye movements track unheard acoustic speech

Neural speech tracking while observing silent speakers has been repeatedly demonstrated by us (Hauswald et al., 2018; Suess et al., 2022) and other groups (Aller et al., 2022; Bourguignon et al., 2020). In recent research (Gehmacher et al., 2024; Schubert et al., 2023) involving naturalistic speech, we found that eye movements track the acoustic speech envelope, especially when attended (see Jin et al., (2018). for artificially rhythmic speech). Here, using EOG estimates instead of eye tracking, we studied ocular unheard speech tracking during the perception of silent lip movements. Indeed, we demonstrate that processing silent lip movements leads to ocular tracking of the unheard speech envelope, which occurs in a frequency range below 1 Hz. We show that this tracking is not trivially explained by the visual processing of the speaker’s lip movements, as the forward versus backward effect is not observed when using the lip movements signal for coherence analysis. These results suggest a learned connection between visual stimuli and auditory speech, expressed in terms of eye movement behavior.

Consistent with this, we found a significant effect for the acquired DHH group, in the same frequency range as in the hearing group. However, this effect was not observed in the congenitally deaf group. When testing for differences between the three groups in terms of ocular unheard speech tracking, the acquired DHH group showed increased tracking compared to the other groups when considering both lip- and speech-tracking.

As Gehmacher et al., (2024) show that ocular tracking is related to attention, the acquired DHH group may focus more on the lip movements in the forward condition as they attempt to make sense of it. This might be because, on one hand, they had some hearing experience, but on the other hand, they had to rely on visual input once they became deaf. This could make them particularly adept at extracting relevant linguistic features from lip movements, underscoring the importance of ocular speech tracking for audiovisual integration and understanding.

The hypothesis that eye movements underlie a learned association between auditory and visual speech may explain the absence of effects in congenitally deaf participants. However, we cannot conclusively determine the absence of speech tracking in congenitally deaf individuals, as the small sample size in this group does not provide sufficient statistical power for definitive conclusions. Additionally, it remains uncertain whether the participants in the congenitally deaf group were genuinely born deaf or experienced significant hearing impairment from birth, which could mean they acquired some audiovisual speech properties. Research demonstrating ocular speech tracking (Gehmacher et al., 2024; Schubert et al., 2023), sound-elicited movements in mice (Bimbard et al., 2023), and eye movements involved in attending to and listening to basic auditory tones (Popov et al., 2021), combined with this investigation, offer a new perspective on (eye) movements as crucially involved in auditory processing and attention. As in this dataset not for all participants EOG was provided, we used the vertical ICA-eye component. Importantly, for the participants with clean EOGs, we found the same effects as using the ICA-eye component.

### Low-frequency cortical tracking of unheard speech in temporal and central regions

Testing for differences between the forward vs. backward condition in whole-brain speech coherence revealed the most pronounced effects in auditory such as in inferior motor and somatosensory areas in low delta frequencies (≤1.3 Hz). The area of the inferior motor and sensory cortex is involved in lip movements, such as in speech production (Kern et al., 2019). For silent lip-reading, this location is especially interesting, as the discrimination certain speech sounds (e.g., ‘ba’ and ‘da’) are articulator-specific (meaning that the lip movements help to discriminate those), and are impaired when the lip area of the inferior primary motor cortex is disrupted with TMS (Möttönen et al., 2013, 2014; Möttönen & Watkins, 2009). The lip area is located in the inferior precentral gyrus, where we observe strong forward vs. backward differences in this study. Furthermore, the subcentral gyrus is involved in spech-related movements and human speech production (Eichert et al., 2021). Evidence for silent speech tracking is quite established in the occipital/visual cortices (Aller et al., 2022; Bröhl et al., 2022; Hauswald et al., 2018; Suess et al., 2022).

However, in the present study, using a data-driven whole-brain analysis, we do not find those occipital effects. Within a region of interest, averaging over frequencies <1 Hz, there is a tendency towards higher forward speech tracking in the occipital area (Fig. 2C). In line with Bourguignon et al., (2020), we report silent speech tracking effects in low-frequency bands <1 Hz in temporal regions. Likewise, Aller et al., (2022) show that in silent lip-reading, the auditory cortices can restore auditory information from visual information when no auditory stimulation is present. Bröhl et al., (2022) also show that lip-reading performance is related to the tracking of the unheard speech envelope (<1 Hz and 1-3 Hz) in auditory, but not in visual cortices.

Given the above, speech tracking while watching silent speakers might be tracked more occipitally in higher frequencies (Aller et al., 2022; Hauswald et al., 2018; Suess et al., 2022) and seems to be more pronounced in auditory and motor areas <1 Hz (Bourguignon et al., 2020; Bröhl et al., 2022). However, investigations that found effects of higher frequencies (4-7 Hz, Hauswald et al., 2018; 1-3 Hz, Suess et al., 2022; 2-6 Hz, Aller et al., 2022; 0.5-3 Hz, Bröhl et al., 2022 (for Pitch)) used higher high-pass filters and shorter epochs compared to our analysis. This can significantly affect the shape of the power and coherence spectra (Schmidt et al., 2023). Using lower high-pass filters and longer epochs, low (<1 Hz) frequency effects have been found in speech tracking (Chalas et al., 2023; Schmidt et al., 2023) and in silent speech tracking (Bourguignon et al., 2020).In auditory speech perception, speech tracking mainly occurs in theta and delta frequencies, while delta (0.3-3 Hz, especially ∼0.6 Hz) is primarily involved in segmenting speech without periodic activity based on speech onsets, such as the beginning of a sentence (Chalas et al., 2023). In this study, the sentence rate is ∼0.25 Hz, so low-frequency tracking can help parse the sentences. Silent lip-reading studies have also shown that from lip movements, mainly slower, delta auditory information is reflected in the brain (Bourguignon et al., 2020). Delta speech tracking peaks <1 Hz (Chalas et al., 2023), and blinks also occur with a frequency below 1 Hz (Jin et al., 2018).

### Ocular speech tracking is not directly related to cortical speech tracking

Our findings indicated that the ocular and neural speech tracking exhibited highly overlapping coherence spectra (Fig. 1B & C). To investigate whether cortical speech tracking is driven by ocular tracking, we calculated partial coherence between neural activity and the unheard speech envelope and partialized out the ocular activity. The results demonstrated that the cortical tracking of the unheard speech envelope is not directly influenced by ocular tracking. Nevertheless, this raises the question of how two highly similar processes here (see Fig. 1A) can occur in the brain and in the eyes, but be fully independent, and how the eyes can reflect the learned connection between auditory and visual speech. Possibly, both the ocular and the cortical tracking of the unheard speech is triggered by a common source that is responsible for the transformation from the visible lip movements into the associated but unheard speech signal. In recent research, investigating ocular tracking in auditory speech using mTRF and a moderation analysis to control for eye movements, Gehmacher et al., (2024) showed that eye movements drive some of the cortical tracking (for replication see Schubert et al., 2023). Also in mice, sounds elicited movements and visual brain activity, and the movements were enough to explain visual brain activity (Bimbard et al., 2023). However, there are two main differences to this data 1) we do not have auditory signals, only visual input. In the other case, the information comes from the auditory system. 2) The cortical effects explained by the eye movements were visual (Bimbard et al., 2023). In this data, the effects appear more temporally including motor cortices. To sum up, here the evidence is clear, that the ocular speech tracking effects do not explain the cortical speech tracking, as the partial coherence did not change the results.

### Limitations and future implications

It is very unlikely, that the eyes are reflecting independent processes from neural activity. In this study, partial coherence could not serve to reveal the connection between both. In future investigations, we aim to test more methods, that can also give us time-resolved insights into the connection between ocular and neural unheard speech tracking. Furthermore, we do not have eye-tracking data in this dataset. Using EOG-ICA components instead is a limited measure, and eye tracking would make a stronger point for our hypothesis. Contrary to eye-tracking, which measures only eye movements, EOG electrodes might also capture some brain data. Additionally, it might be that some effects remain undetected, eye-tracking provides more precise information about the direction and the extent of the movements. Using eye-tracking, it could be also investigated whether the focus is more on the lips or on the eyes of the speaker and whether this varies between different groups. In this work, instead of the EOG only the vertical blink-related ICA eye component was analyzed because it was most consistently present over all subjects. Even though the ICA eye component is statistically independent from the other ICA components, we cannot exclude brain data with certainty. Despite this, using the EOG or the ICA eye component, interesting questions such as ocular tracking in blind subjects (who also might not have eyes) or subjects with closed eyes can be addressed. This cannot be achieved using eye-tracking which makes this approach an important addition, which is also emphasized by our strong tracking effects found in the EOG data. New approaches enabling saccadic detection (Madariaga et al., 2023) can improve this method and promise more insights in ocular speech tracking in future research. But what can be said here, is that the eye movements do play a role in speech tracking. This opens a new perspective in language processing research and emphasizes how highly integrated the human senses are. Also, when eye tracking is possible, in future studies both measures can be integrated and compared to make a stronger point.

## Conclusion

The eyes are often referred to as windows of the mind, and it has been shown that the eyes are engaged in many processes beyond just visual ones (Van Gompel, 2007). Recently, it has also emerged that the eyes are involved in attending to and listening to basic auditory tones (Popov et al., 2021) and complex, learned language (Gehmacher et al., 2024; Schubert et al., 2023). This highlights the general role of eye movements in speech perception and offers a new interpretation of multimodality in speech perception. Here, we emphasize the relevance and involvement of eye movements in speech perception. We demonstrate ocular unheard speech tracking while observing silent lip movements on the one hand, and we also suggest mechanisms in ocular and cortical speech tracking that are not directly influenced by each other but might be controlled by a common source. Furthermore, the ocular speech tracking effects were only revealed in individuals with hearing experience, suggesting that audio-visual listening experience is necessary for ocular speech tracking in silence. The absence of differences in the congenitally deaf group, combined with the enhanced effects in the acquired DHH group, suggests that these effects only exist in individuals with hearing experience encompassing the critical period for verbal speech acquisition. Overall, this study provides insights into the role of eye movements in speech processing and raises important questions about if and how the eyes are functionally involved in neural speech processing.

## 4. Methods

### 4.1 Participants

Originally, 75 individuals (31 male, 44 female, 3 left-handed, mean age: 42, SD: 14) participated in this study. Forty-nine hearing participants were re-analyzed from the study by Suess et al., (2022), and 26 hearing-impaired participants were recruited additionally. Nineteen of them acquired DHH after birth and had various diagnoses concerning their listening ability, such as bilateral severe hearing loss, bilateral profound hearing loss, and single-sided deafness. Seven participants were born deaf (see Suppl. Mat.). This led to three groups: the hearing group, the DHH group, and the congenitally deaf group. One of the subjects from Suess et al. was excluded, as no ICA-eye component could be identified.

Initially, the intention was to categorize the deaf participants into pre-lingually (<3 years) and post-lingually (>3 years) deafened groups. However, participants who had audiovisual language exposure after birth emerged as significant outliers, exhibiting patterns more akin to those of the post-lingually deafened group. Consequently, we re-categorized the participants into new groups: those with audiovisual language experience and those without. With respect to development, newborns exhibit neural speech tracking (Florea et al., 2024), and fetuses demonstrate differential reactions to familiar versus unfamiliar languages (Kisilevsky et al., 2009; Minai et al., 2017). Therefore, this early (<3 years) audiovisual language experience might be sufficient to elicit low-frequency speech tracking during silent lip-reading. Requirements for participation included normal or corrected-to-normal vision, no prior history of neurological or psychological disorders, no intake of any medication or substance that could influence the nervous system, and no ferromagnetic metal in the body. The experimental procedure was approved by the University of Salzburg ethics committee and was carried out in accordance with the Declaration of Helsinki. Participants signed consent forms.

The criteria for the hearing group included normal hearing and German as a mother tongue. For the congenitally deaf group, the criteria were being born deaf and having sign language as a native language. The acquired DHH group consisted of individuals who lost their hearing after acquiring their native language, German. The congenitally deaf subjects were recruited from local deaf associations, while the acquired DHH group came to the university hospital to receive cochlear implants and participated in our study before their implantation. As compensation, participants received 10€ per hour or course credits.

### 4.2 Procedure

The procedure was identical to the one used in Suess et al., (2022). Participants were instructed to pay attention to the lip movements of the speakers and passively watch the mute videos. They were presented with 6 blocks of videos, and in each block, 2 forward and 2 backward videos were presented in random order. The experiment lasted about an hour including preparation. The experimental procedure was programmed in Matlab with the Psychtoolbox-3 (Brainard, 1997) and an additional class-based abstraction layer (https://gitlab.com/thht/o_ptb) programmed on top of the Psychtoolbox (Hartmann & Weisz, 2020).

### 4.3 Data acquisition

Before the MEG recording, five head position indicator (HPI) coils were applied to the subjects’ scalps. Anatomical landmarks (nasion and left/right pre-auricular points), the HPI locations, and approximately 300 headshape points were sampled using a Polhemus FASTTRAK digitizer. Auditory stimuli were presented binaurally using MEG-compatible pneumatic in-ear headphones (SOUNDPixx, VPixx Technologies, Canada). For recording neural activity, a whole-head 306-sensor MEG system (Elekta Neuromag Triux, Elekta Oy, Finland) in a magnetically shielded room (AK3b, Vacuumschmelze, Germany) was used. Frequencies in the range of 0.1–330 Hz were recorded at a sampling rate of 1 kHz. The head position inside the MEG helmet was continuously monitored during the experiment using five head-tracking coils. The coils indicating the head position, three anatomical fiducials, and at least 150 individual head-surface points on the scalp and the nose were localized in a common coordinate system with an electromagnetic tracker. As a standard procedure in the lab, we also measured EOG and electrocardiogram (ECG).

A signal space separation (SSS) algorithm (Taulu et al., 2004; Taulu & Simola, 2006), implemented in MaxFilter version 2.2.15 provided by the MEG manufacturer, was used. The algorithm removes external noise from the MEG signal (mainly 16.6 Hz and 50 Hz, plus harmonics) and realigns the data to a common standard head position ([0 0 40] mm, -trans default MaxFilter parameter) across different blocks, based on the measured head position at the beginning of each block.

### 4.4. Stimuli & Extraction of stimulus features (Suess et al., 2022)

Videos were recorded with a digital camera (Sony NEX FS100) at a rate of 50 frames per second, and the corresponding audio files were recorded at a sampling rate of 48 kHz. The videos were spoken by two female native German speakers. The speakers were told to narrate the text with as little additional face and body expressions as possible to avoid influences from other facial gestures (as our main interest was the processing of the lip movements). For both speakers the syllable rate was 3.3Hz (calculated with syllable nuclei (de Jong & Wempe, 2007) computed using the Python library *parselmouth* (Jadoul et al., 2018)). The sentence rate was ∼ 0.25 Hz. This was calculated manually dividing the number of sentences by the time. We uploaded two example videos on our OSF-page (https://osf.io/ndvf6/). One speaker was then randomly chosen per subject and kept throughout the experiment, so that each participant only saw one speaker. The stimuli were taken from the book “Das Wunder von Bern” (“The Miracle of Bern”; https://www.aktion-mensch.de/inklusion/bildung/bestellservice/materialsuche/detail?id=62), which was delivered in an easy language. The easy language does not include any foreign words, has a coherent verbal structure, and is easy to understand. We used simple language to avoid limited linguistic knowledge from interfering with possible lip-reading abilities. Twenty-four pieces of text were chosen from the book and recorded by each speaker, lasting between 33 and 62 s, thus resulting in 24 videos. Additionally, all videos were reversed, which resulted in 24 forward videos and 24 corresponding backward videos. Forward and backward audio files were extracted from the videos and used for the data analysis. Half of the videos were randomly selected to be presented forward and the remaining half to be presented backward. The videos were backprojected on a translucent screen in the center of the screen by a Propixx DLP projector (VPixx Technologies, Canada) with a refresh rate of 120 Hz per second and a screen resolution of 1920 x 1080 pixels. The translucent screen was placed approximately 110 cm in front of the participant and had a screen diagonal of 74 cm.

The lip movements of every speaker were extracted from the videos with a MATLAB script (Park et al., 2016; Suess et al., 2022), where the lip contour, the area, and the horizontal and vertical axis were calculated. Only the area was used for the analysis, which leads to results comparable to using the vertical axis (Park et al., 2016). The lip area signal was up sampled from 50 Hz to 150 Hz using FFT-based interpolation, to match the MEG data for further analysis.

The acoustic speech envelope was extracted with the Chimera toolbox from the audio files corresponding to the videos, which constructed nine frequency bands in the range of 100 – 10000 Hz as equidistant on the cochlear map (Smith et al., 2002). The respective cut-off values for the nine frequency bands were as follows: 101, 220, 402, 680, 1103, 1748, 2732, 4231, 6517 and 10000. Those values are based on the cochlear frequency maps for the cat scaled to fit the human frequency range of the hearing (Liberman, 1982). Then the sound stimuli were band-pass–filtered in these bands with a fourth-order Butterworth filter to avoid edge artifacts. For each of the frequency bands, the envelopes were calculated as absolute values of the Hilbert transform and then averaged to get the full-band envelope for coherence analysis (Gross et al., 2013; Keitel et al., 2017). This envelope was then down sampled to 150 Hz to match the preprocessed MEG signal.

### 4.5 Preprocessing

Data preprocessing was done in MNE–Python (Gramfort, 2013).With Independent Component Analysis (fast ICA; Hyvarinen, 1999), after applying a 1 Hz high-pass filter, 50 linearly mixed sources were separated, and heartbeat, eye-blink, and eye movement artifacts were selected from one example subject. Using template matching (Campos Viola et al., 2009), these components were found for all subjects and were rejected from the MEG data. As the data was originally not collected to investigate eye movements, out of 73 subjects, only 38 had eye-tracking data included, and 45 had EOG data of good quality. In order to analyze the full sample, we used the ICA eye component that showed the highest correlation with the vertical EOG in one subject to detect the relevant components for the other subjects using template matching (Campos Viola et al., 2009). This component was used as it could be identified most consistently over subjects, and also Jin et al., (2018) found their main effects in the vertical EOG. The ICA-eye component is further referred to as ocular activity.

In the previous analysis by Suess et al., (2022), a 1 Hz high-pass filter was used. As (Bourguignon et al., 2020) found the main effect of silent speech tracking below 1 Hz, and eye movements also have slow components, we decided on a lower frequency range from 0.1–5 Hz for the coherence analysis. Therefore, MEG data, ocular activity, as well as the speech envelope and the lip movements, were filtered with a high-pass filter of 0.1 Hz and a low-pass filter of 12 Hz using overlap-add FIR filtering with symmetric linear FIR filters and a Hamming window. Filter length was based on the transition regions (6.6 times the reciprocal of the shortest transition band). For the coherence analysis, the data was segmented into epochs of 6 seconds to ensure sufficient resolution of the low-frequency oscillations. Each block was assigned to one of the two conditions.

### 4.6 Source projection of MEG data

Source projection of the epoched data was done with MNE-Python (Gramfort, 2013). A semi-automatic co-registration pipeline was used to coregister the FreeSurfer “fsaverage” template brain (Fischl, 2012) to each participant’s head shape. After an initial fit using the three fiducial landmarks, the co-registration was refined with the Iterative Closest Point (ICP) algorithm (Besl & McKay, 1992). Head shape points that were more than 5 mm away from the scalp were automatically omitted. The subsequent final fit was visually inspected to confirm its accuracy. This semi-automatic approach performs comparably to manual coregistration pipelines (Houck & Claus, 2020). A single-layer boundary element model (BEM; (Akalin-Acar & Gençer, 2004) was computed to create a boundary element method (BEM) solution for the “fsaverage” template brain. Next, a volumetric source space was defined, containing a total of 5124 sources. Subsequently, the forward operator (i.e. lead field matrix) was computed using the individual coregistrations, the BEM, and the volume source space. Afterwards, the data were projected to the defined sources using the linearly constrained minimum variance (LCMV) beamformer method.

### 4.7. Coherence calculation

Coherence was calculated in Fieldtrip (Oostenveld et al., 2011). Therefore, the epoched data in source space was imported from MNE. For the single epochs, we conducted a multitaper time-frequency analysis with multiple tapers based on discrete prolate spheroidal sequences (DPSS). The amount of spectral smoothing through multi-tapering was set to 0.5. Frequencies of interest were from 0.166 Hz to 5 Hz in steps of 0.166. The complex Fourier spectrum was then used to calculate the coherence.

In the first step, only the coherence between the stimuli (lip movements and speech envelope) and the eye movements (ICA-eye components) was calculated. In the second step, the coherence between the stimuli (lip movements and speech envelope) and each voxel of the brain data was calculated. In a third step, in order to gain further insights into the role of eye movements in speech-brain coherence, partial coherence was calculated between stimuli (lip movements and speech envelope) and each voxel of the brain data, with eye movements (ICA-eye components) partialized out. This means that only the coherence between the brain and stimuli data that cannot be attributed to eye movements is calculated.

### 4.8. Statistics

To test for possible differences between the forward vs. backward condition, a cluster permutation dependent t-test was calculated (Maris & Oostenveld, 2007). The cluster permutation t-test controls for multiple comparisons (frequency steps, voxels). For the two-sided t-test, the p-value threshold for the clusters was set to 0.01. This analysis was conducted for the results in the frequency domain. To obtain results in source space, the cluster permutation was run over all voxels and over the frequencies. To test for potential differences between the three groups, we first substracted the coherence of the backward trials from the forward coherence for both, lip and speech coherence. Then we ran a cluster permutation 2×3 ANOVA on a whole brain level and post hoc tests. The cluster permutation tests were ran in eelbrain (Brodbeck et al., 2021). For the eyes and the single voxel, also a 2×3 ANOVA and post hoc tests were run in pingouin (Vallat, 2018).

## Acknowledgements

Thanks to the whole research team for their support in all the challenges we tackled. Special thanks go to Verena Zehntner and Jessica Deprieux, who recorded the videos, helped with the measurements, and recruited participants. The measurements were mainly run by Manfred Seifter—immense thanks for that! Thomas Hartmann gave technical support and Patrick Reisinger shared analysis scripts. Most grateful acknowledgments go to Nina Suess, who planned and coordinated the experiment and shared the data.

## Data Availability

Find the code on gitlab: https://gitlab.com/KajaRosa/eog_analysis_clean/

## Author contributions

N.S. and A.H. designed the experiment, K.B. analyzed the data, generated the figures, and wrote the manuscript. N.S. recruited participants, A.H. and Q.G. supported the data analysis, and edited the manuscript. G.D. supported to develop the source-reconstruction pipeline. F.S. supported the data analysis. G.H. and S.R. recruited the deaf participants and edited the manuscript. N.W. designed the experiment, acquired the funding, supervised the project, and edited the manuscript.

## Supplemental information

**Supp. 1:**
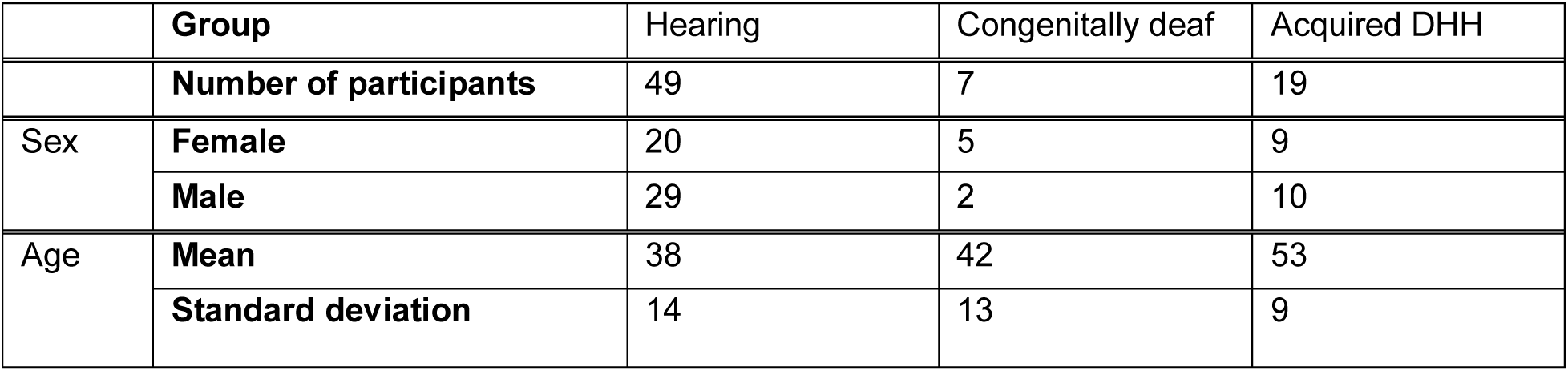
Group information

**Supp. 2:**
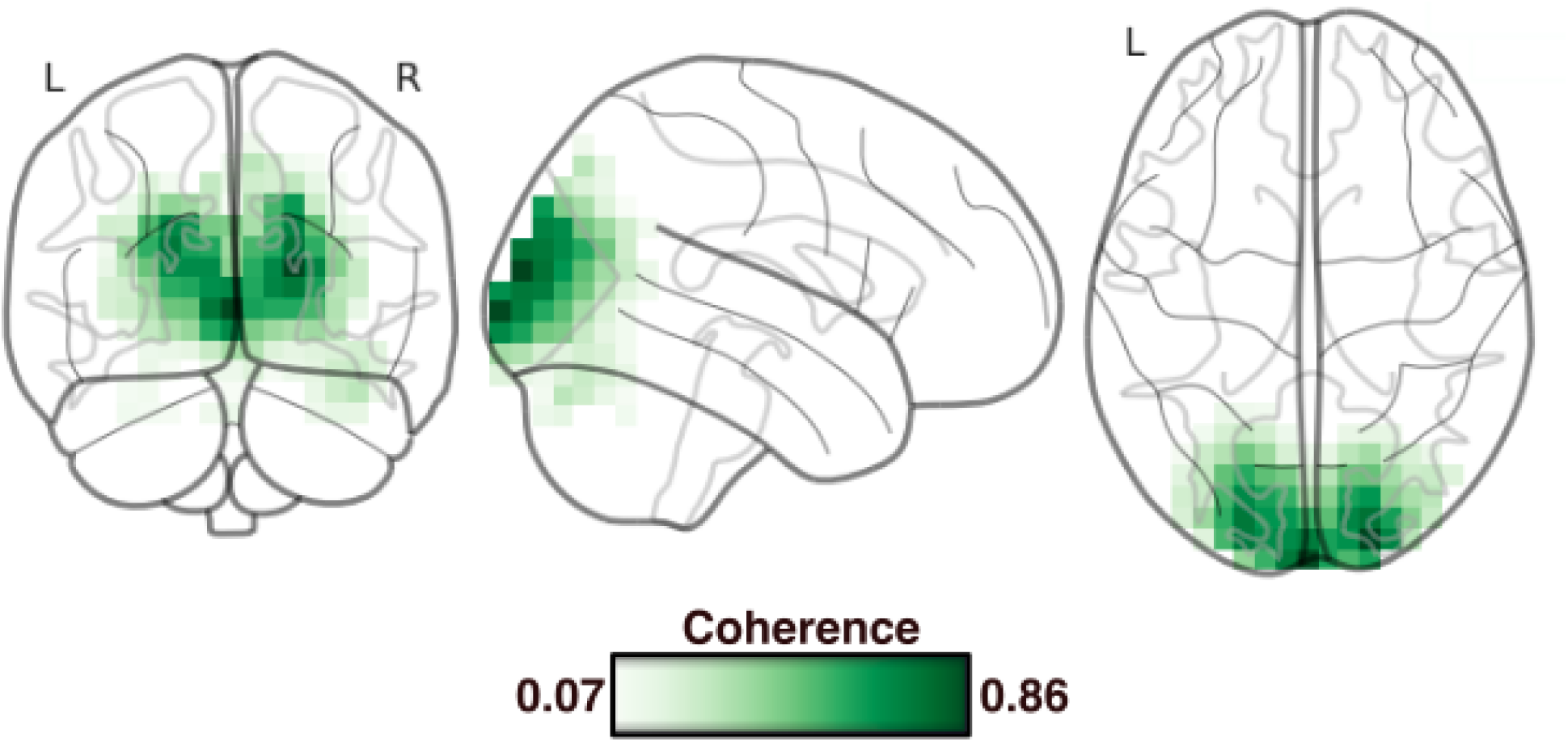
Sanity check for the source reconstruction: Voxel with the strongest lip coherence

**Supp. 3:**
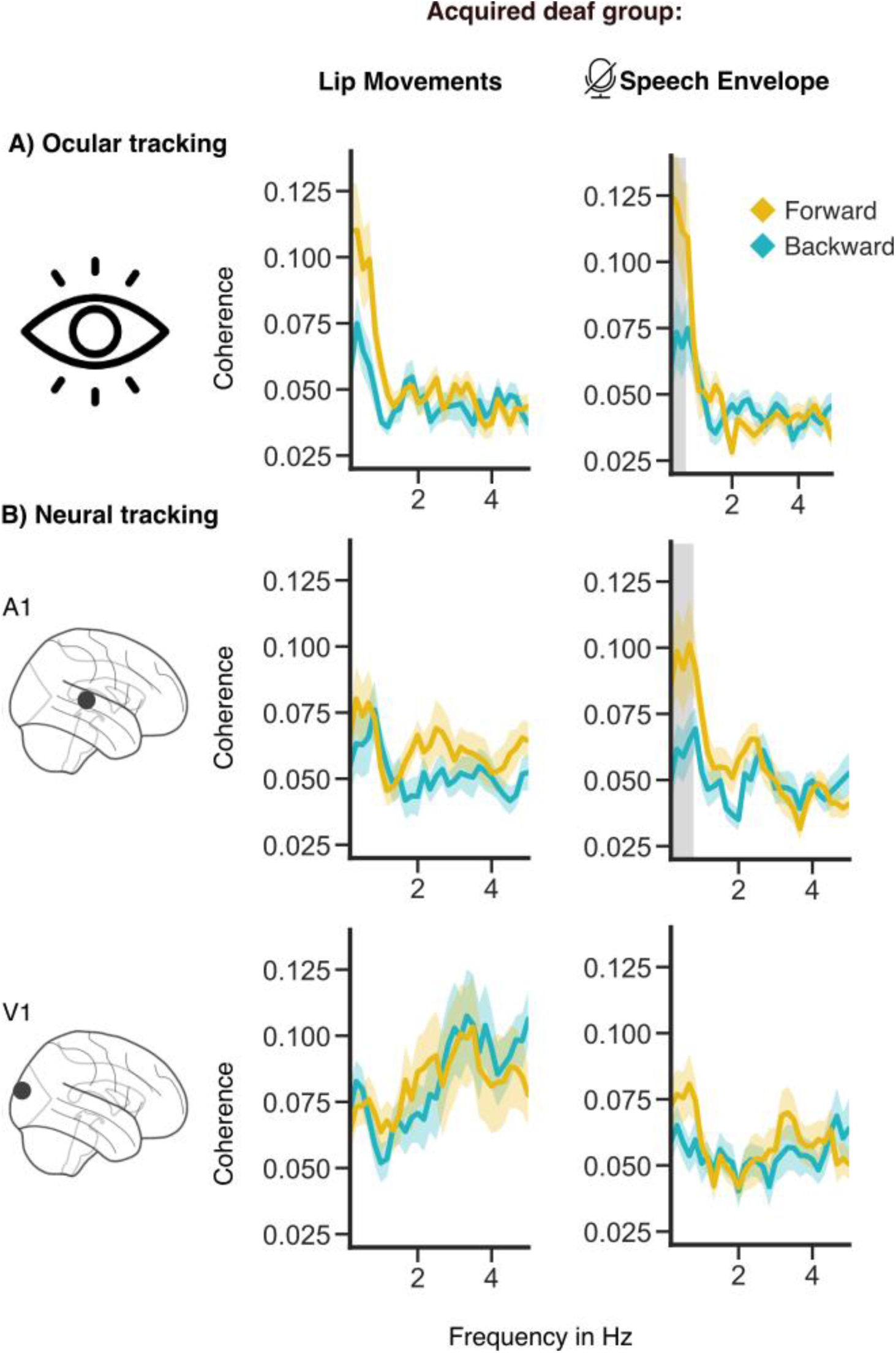
Effects for the acquired DHH group A) results of Coherence calculation of speech envelope and the lip movements with the selected ICA eye component. B) Coherence calculation at the strongest voxel in iii) in the primary auditory and visual cortex. Significant clusters are marked in gray (N=19).

**Supp. 4:**
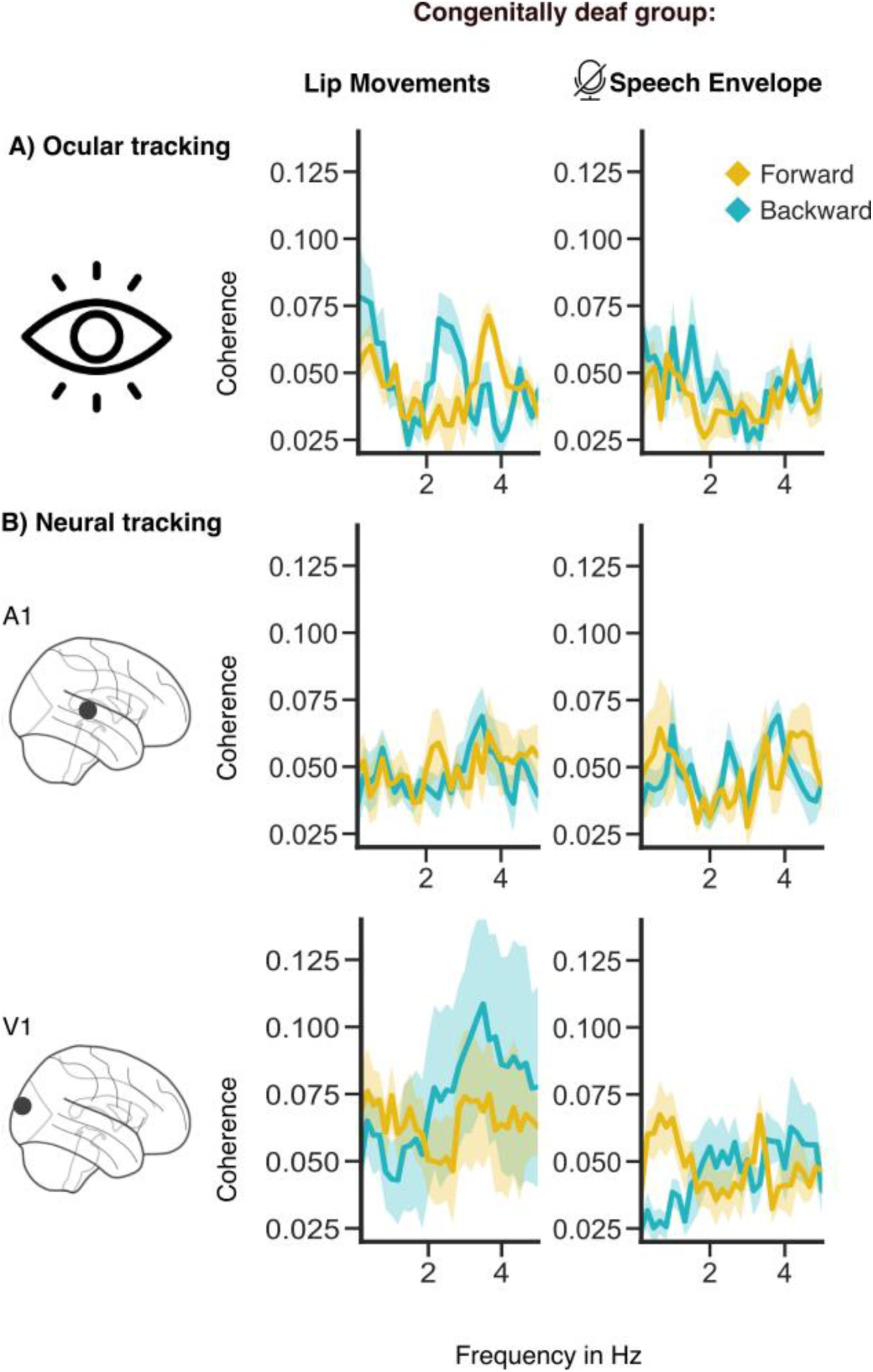
Effects for the acquired DHH group A) results of Coherence calculation of speech envelope and the lip movements with the selected ICA eye component. B) Coherence calculation at the strongest voxel in iii) in the primary auditory and visual cortex. Significant clusters are marked in gray (N=7).

